# Engineering Translational Resource Allocation Controllers: Mechanistic Models, Design Guidelines, and Potential Biological Implementations

**DOI:** 10.1101/248948

**Authors:** Alexander P.S. Darlington, Juhyun Kim, José I. Jiménez, Declan G. Bates

**Affiliations:** Warwick Integrative Synthetic Biology Centre, School of Engineering, University of Warwick, Coventry, CV4 7AL, UK.; Faculty of Health and Medical Sciences, University of Surrey, Guildford, GU2 7XH, UK.

## Abstract

The use of orthogonal ribosomes in combination with dynamic resource allocation controllers is a promising approach for relieving the negative effects of cellular resource limitations on the modularity of synthetic gene circuits. Here, we develop a detailed mechanistic model of gene expression and resource allocation, which when simplified to a tractable level of complexity, allows the rational design of translational resource allocation controllers. Analysis of this model reveals a fundamental design trade-off; that reducing coupling acts to decrease gene expression. Through a sensitivity analysis of the experimentally tuneable controller parameters, we identify how each controller design parameter affects the overall closed-loop behaviour of the system, leading to a detailed set of design guidelines for optimally managing this trade-off. Based on our designs, we evaluated a number of alternative potential experimental implementations of the proposed system using commonly available biological components. Finally, we show that the controller is capable of dynamically allocating ribosomes as needed to restore modularity in a number of more complex synthetic circuits, such as the repressilator, and activation cascades composed of multiple interacting modules.

## 1 Introduction

Ensuring circuit modularity, i.e the independent and predictable functioning of different circuit processes, remains a key goal in synthetic biology. If modules are independent then they can be recombined to produce novel functions which can be predicted from previous characterisation. This approach is commonly used in electronics and computer science where complex functions are broken down into independent modules, which can be assembled to form new systems.

However, in synthetic circuits, there is often a failure of modularity, with gene circuits based on purportedly well characterised components needing iterative rounds of redesign and re-experimentation to obtain functional implementations. Modularity fails for a variety of reasons: (i) unexpected cross talk between modules due to component re-use [1], (ii) subtle changes in gene regulation due to unforeseen effects of combining DNA sequences [2], (iii) retroactivity effects where the titration of a transcription factor to a downstream module affects the behaviour of the upstream module [3], and, (iv) the use of a common limited pool of resources for gene expression [4]. Careful selection of components can ameliorate the effects of (i) by ensuring modules do not have off target effects [5]. The introduction of ‘insulator elements’ such as ribozymes can reduce the effects of (ii) [6] and the development of buffer circuits allows loading fracture due to retroactivity (iii) to be reduced [7]. The question of how to optimally manage the effects of cellular resource limitations on circuit modularity, however, remains an open problem.

During exponential growth, the number of RNA polymerases (RNAP) and ribosomes in the cell remains constant. This results in a fixed pool of gene expression resources. Whilst both resources are finite, numerous experimental studies have shown it is the number of free ribosomes in system which is the main limitation on gene expression [8, 9, 10, 11, 12]. The sharing of this fixed resource across genes leads to a phenomenon known as gene-coupling. This results in the emergence of non-regulatory interactions between co-expressed genes [4], since each synthetic circuit module will utilise as many ribosomes as possible at any one moment, as determined by parameters such as mRNA levels or RBS strength.

To illustrate the problem mathematically, consider the number of free ribosomes as a function of the ribosome supply and demand as

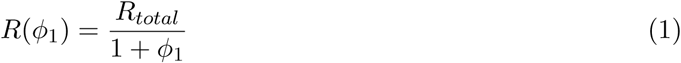

where *R*_*Total*_ is the total number of ribosomes available and *ϕ* is the demand, [10, 4]. Upon the addition of another demand *ϕ*_2_, the free ribosome number becomes:

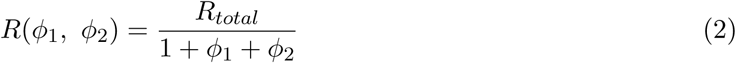

As *ϕ* > 0 in all cases, by definition *R*(*ϕ*_1_, *ϕ*_2_) < *R*(*ϕ*_1_). This decrease in free ribosome number reduces the rate of downstream processes, such as mRNA-ribosome binding, as a consequence of the law of mass action. This leads to a decrease in other modules as a new module is induced (this is often termed coupling [13, 14]): activation of one circuit module effectively inhibits other modules. In this case the supply of ribosomes (the numerator) is determined by the cell and supply is constant regardless of the demand (the denominator) - i.e. there is no control of *R*_*Total*_.

To mitigate this decrease in free ribosomes upon the addition of new genes, consider a system where the supply of total ribosomes can be matched to the circuit’s demand for ribosomes. Let this malleable ribosome pool be *R*′:

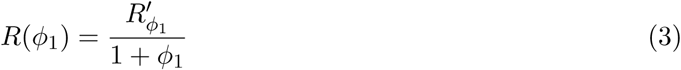

Upon the addition of another gene the demand (the denominator) increases as before:

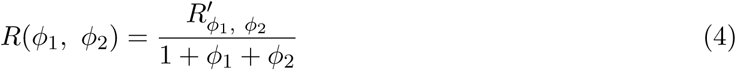

However, in this ideal system we can now increase the supply of *R*′ (i.e. increase the numerator) to make *R*(*ϕ*_1_) = *R*(*ϕ*_1_, *ϕ*_2_. This maintains the free ribosome pool available to the circuit and so removes ribosome-mediated gene coupling.

In a previous work, we have experimentally realised a prototype of a feedback controller that can dynamically allocate more ribosomes to the circuit when required [14] (Figure 1a). Increasing total ribosome number is not biologically feasible, so our controller acts to dynamically allocate the translational capacity *between* host and circuit genes in response to circuit demand, thus relieving the effects of resource limitations on the circuit [14]. This is achieved by regulating the production of a pool of quasi-orthogonal ribosomes. These specialised circuit-specific ribosomes can be created by expressing an orthogonal 16S rRNA and replacement of the natural ribosome binding site in circuit genes, or other genes of interest, with complimentary synthetic ribosome binding sites. The o-16S rRNA replaces the endogenous host 16S rRNA in a fraction of the host ribosomes creating a separate translational resource which is targeted to circuit genes by binding the complimentary synthetic RBS sequence. By placing the production of the o-16S rRNA under the control of a constitutively expressed repressive transcription factor which itself uses the orthogonal ribosome pool for its own translation, we created a feedback controller which produces o-16S rRNA, and hence orthogonal ribosomes, in response to circuit demand. As circuit genes are induced, they sequester o-ribosomes for their own expression resulting in a fall in the expression of the regulator. This relieves the repression of the o-16S rRNA, resulting in increased o-16S rRNA production and increased o-ribosome co-option. Thus the controller implements a negative feedback loop. See Figure S1 for a schematic of the biological implementation of this feedback loop. Of course, the controller cannot mitigate intrinsic limitations arising from the fact that the total number of ribosomes in the cell is finite. Rather, the controller acts to dynamically manage the allocation of translational activity between host and circuit genes in the most efficient way, by increasing circuit capacity as circuit demand requires it.

**Figure 1:**
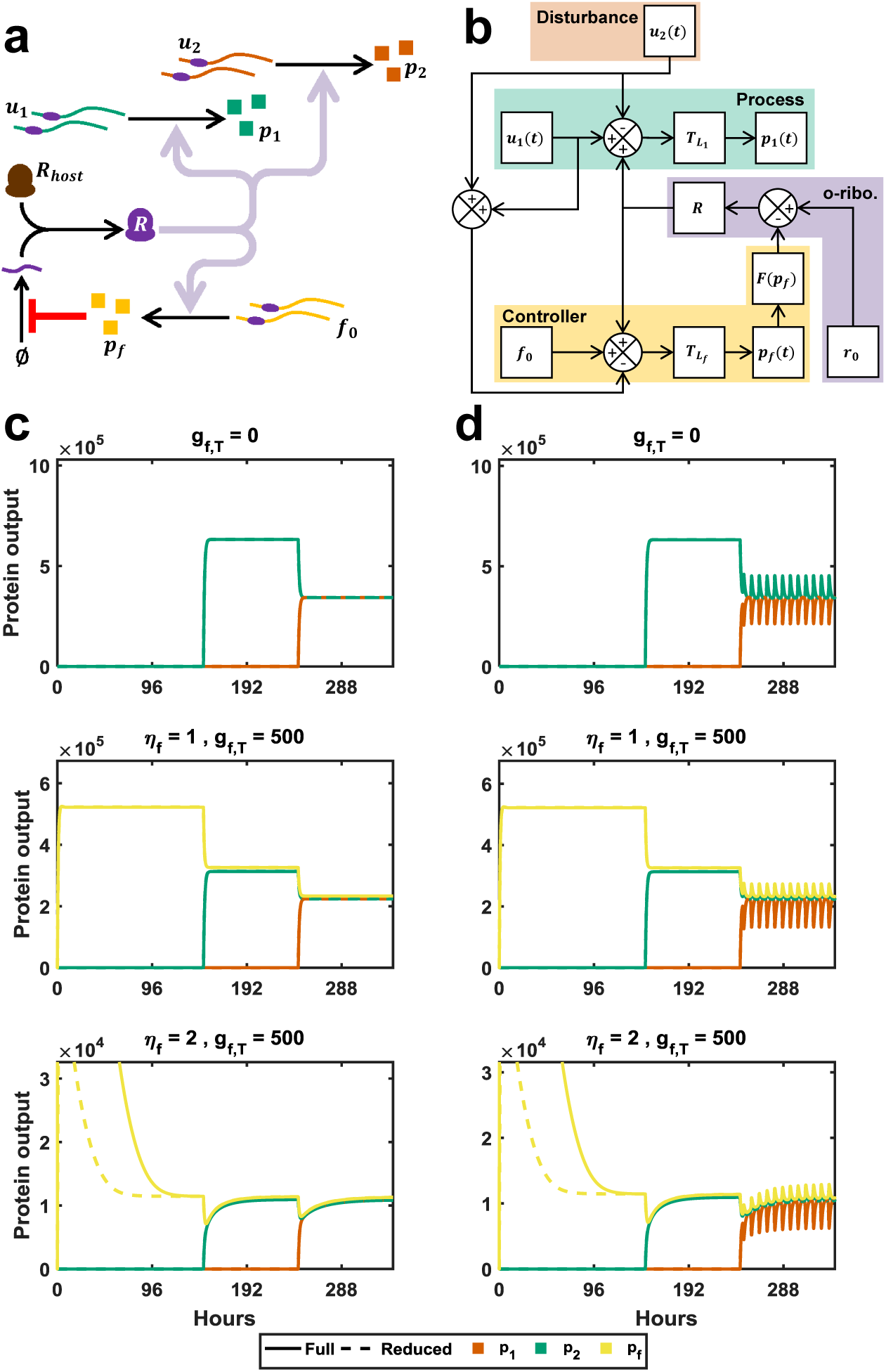
Development of a genetic feedback controller model. **(a)** Schematic of the negative feedback loop implementation. **(b)** Block diagram of the controller. The process, highlighted in green, converts the input *u*_1_ into protein output *p*_1_ utilising the o-ribosome pool *R.* The input into a second process (not shown) *u*_2_ acts as a disturbance to the first process which is ameliorated by the effect of the controller. The controller protein is constitutively expressed (*f*_0_ signal) so the output pf is dependent upon *R.* As inputs *u*_1_ disturb *R* the level of *p*_*f*_ changes (i.e. as *u*_1_ increases, *P*_*f*_ decreases). As *p*_*f*_ is a repressor the disturbance signal is inverted in the *F*(*P*_*f*_) block. **(c)** The reduced model successfully captures the behaviour of the full model. Numerical simulations carried out as described in the Methods. Parameters used are derived from [17]. A range of controllers are shown: open loop with no controller *g*_*f*, *T*_ = 0 nM, a linear controller *η*_*f*_ = 1,*g*_*f*, *T*_ = 500 nM, a non-linear controller *η*_*f*_ = 2,*g*_*f*, *T*_ = 500 nM. Inputs are as follows: *u*_1_ = *u*_2_ = 0 nM except *u*_1_(*t* > 148) = 500 nM and *u*_2_ (*t* > 244) = 500 nM **(d)** As in **(c)** with inputs *u*_1_ = *u*_2_ = 0 except *u*_1_(*t* > 148) = 500(cos(0.8t) + 1) nM and *u*_2_ (*t* > 244) = 500(cos(0.8t) + 1) nM.

In this paper, we develop detailed mechanistic models of the orthogonal translation system that can be used for the purposes of designing optimal resource allocation controllers. Using such models, we demonstrate how improved resource allocation controllers can be rationally designed to decouple the expression of different genes, and develop design rules for how the tuning of different controller design parameters can act to separately specify either the dynamic response time or overall protein output (i.e. gain) of the circuit. Based on these design rules, we identify and evaluate a number of alternative potential experimental implementations of the proposed translational controllers. Finally, we demonstrate the potential of resource allocation controllers to improve the modularity of a variety of complex gene circuits.

## 2 Results and discussion

### 2.1 A mechanistic model of the resource allocation controller

We initially develop a complete mechanistic model of gene expression and the action of the controller, before investigating how this model can be simplified for use as a design tool. We assume that each circuit promoter (*g*_*i*_) can be bound by a multimeric transcription factor (*u*_*i*_) to form a promoter complex (*κ*_*i*_) capable of recruiting a free RNA polymerase (*σ*) to form a translation complex. When transcription occurs, an mRNA (*m*_*i*_) is produced, and the original RNAP polymerase and promoter complex are released. The above interactions are described by the following chemical reactions:

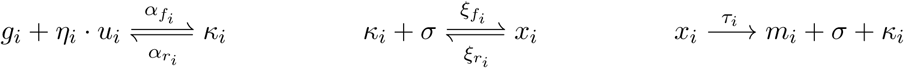

The mRNA is bound by a free [orthogonal] ribosome, *R*, to form a translation complex (*c*_*i*_). Upon translation, a protein (*p*_*i*_) is produced and the mRNA and *R* are released:

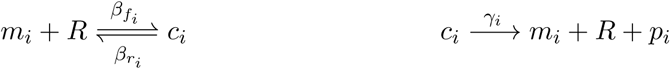

Additionally, both mRNAs and proteins degrade at rates *δ*_*m*_*i*__ and *δ*_*p*_*i*__, respectively. The numbers of promoters for each gene i are conserved such that:

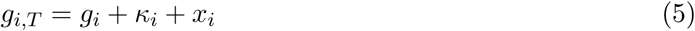

Applying the law of mass action we derive the following ODEs describing the time evolution of the circuit components:

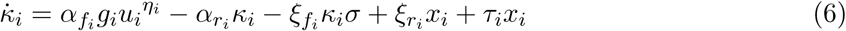

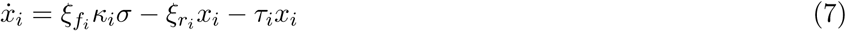

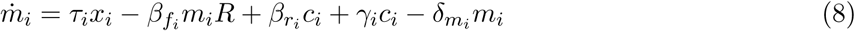

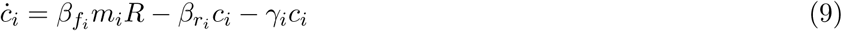

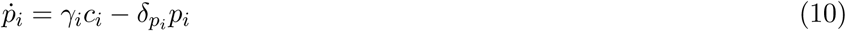

This represents a simple single-input-single-output (SISO) motif and forms the basis of our model. Complex circuits can be constructed by letting the output from one module form the input to another.

To implement our controller we first consider the conversion of host ribosomes (*R*_*host*_) into circuit-specific orthogonal ribosomes (*R*). The orthogonal 16S rRNA gene promoter (*g*_*r*_) recruits *σ* to form a translation complex (*x*_*r*_) which produces the orthogonal rRNA (*r*):

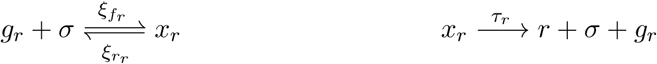

The orthogonal 16S rRNA binds host ribosomes, *R*_*H*_, and so recruits ribosomes to the circuit-only orthogonal pool, *R*:

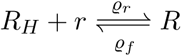

In the presence of the controller the orthogonal rRNA gene is regulated by the repressor *p*_*f*_. The repressor binds the free *g*_*r*_ promoter and prevents the binding of RNA polymerase and associated factors (*σ* in our model):

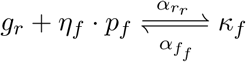

We model expression of the regulator protein by considering the constitutive expression of its mRNA from an unregulated promoter, *g*_*f*_:

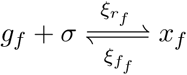

We model the transcription and translation of the repressor’s mRNA and protein in the same manner as the circuit genes, as described above.

The concentration of each promoter is conserved such that:

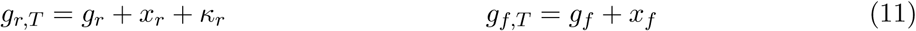

Applying the law of mass action results in the following ODEs describing the production of the repressor and intermediate complexes:

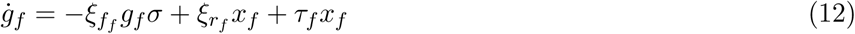

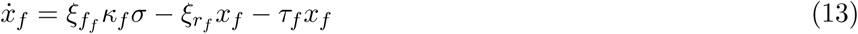

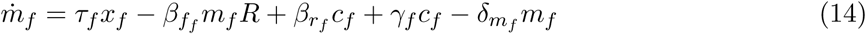

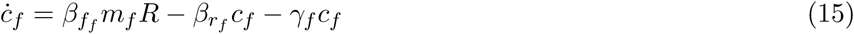

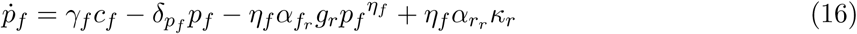

Applying the law of mass action to the o-rRNA promoters and ribosome species yields:

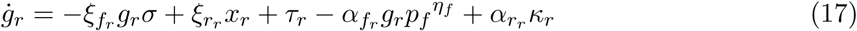

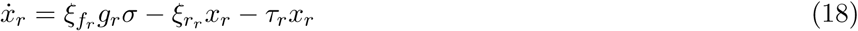

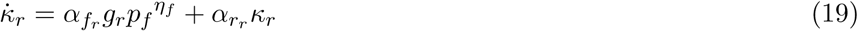

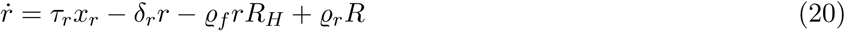

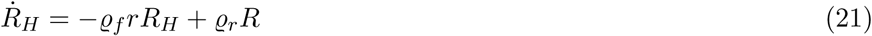

This model is highly complex and contains many forward and reverse reaction rates. Individual binding and unbinding rates for genetic components and proteins are rarely reported in the literature, due to the difficulty in determining their values experimentally because of the effects of confounding factors (such as additional fluxes along the reaction pathway). However, by considering the time-scale separation of the reactions involved, we are able to reduce our model and formulate the forward and reverse reactions in terms of ‘lumped’ dissociation constants. By defining the model in terms of dissociation constants, we are then able to determine suitable genetic components with the desired dynamics from a search of previously published data. The dissociation constants of the RNAP polymerase for the promoter (*k*_*X*_), of the ribosome for the RBS (*k*_*L*_) and of the protein for its binding site (*μ*) are defined as follows:

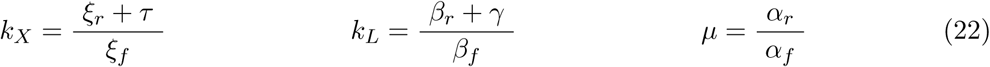

To reduce our model we consider the effect of time-scale separation; different biological processes occur over a range of different time spans with binding/unbinding reactions occurring on the order of milliseconds, transcription and translation taking minutes and protein degradation/dilution occurring over tens of minutes or hours [15]. This effectively separates reactions in time and allows us to apply the assumption that ‘fast’ species, such as RNAs, reach (quasi)-steady state (QSS) instantaneously. By calculating the QSS concentrations of intermediate species and substituting as appropriate we are able to the remove the majority of intermediate gene expression species from our model. We denote the QSS complex of species *y* as *y̅*.

Since current experimental evidence suggests that competition for RNA polymerases does not significantly limit gene expression, we remove RNAP mediated competition by considering each gene to have access to its own small pool of RNA polymerase (e.g. [8, 10]). Additionally, we assume that the dissociation constant for RNA polymerase is much higher than the concentration of free polymerase, consistent with experimental observations [16]. This allows us to reduce the complexity of the expressions by assuming that *σ* + *k*_*X*_ ≈ *k*_*X*_. By applying these assumptions we can simplify Equations 5 to 10 as follows:

The QSS of the activated circuit gene (*x̅*_*i*_) which gives rise to mRNAs is solely a function of the input (*u*_*i*_):

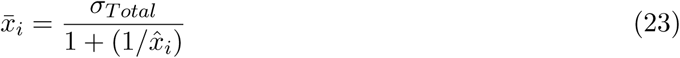

where *x̂*_*i*_ can be considered as a measure of demand for RNAP by gene *i* and is defined as follows:

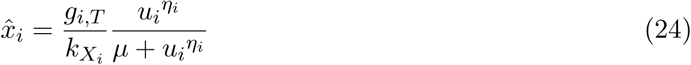

The ODE describing the time-evolution of the protein species is:

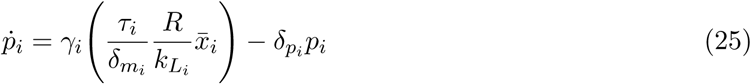

We can also define the constant *ĉ* which is a measure of demand for ribosomes by gene *i*:

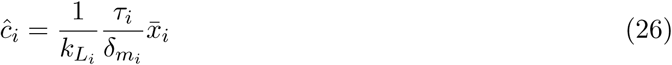

If a single unregulated pool of ribosomes is used for circuit expression then the number of free (host/orthogonal) ribosomes, *R*, available for circuit translation is given by

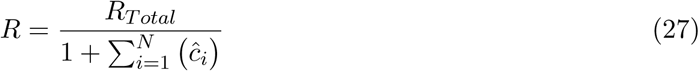

Therefore the response of *p*_1_ depends not only on the input *u*_1_ but also the demand for ribosomes by other genes *ĉ*_*i*, *i*≠1_. This forms the basis of our circuit SISO ‘process’ model (Figure 1b).

Applying the same assumptions to the equations describing the production of the regulator *p*_*f*_ we can reduce equations 11 to 16 to:

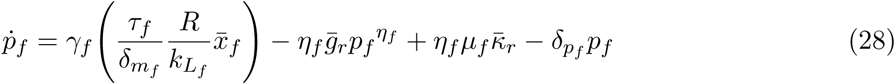

where *x̅*_*f*_ follows the same form as Eq. 23 and *x̂*_*f*_ = *g*_*f*,*T*_/*k*_*X*_*f*__ (*f*_0_ block of Figure 1b). For simplicity we assume that *α*_*f*_*f*__ = 1 and *α*_*f*_*r*__ is equal to the dissociation constant *μ*_*f*_. This forms the basis of the ‘controller module’ shown in Figure 1b, with the action of the controller represented by the *F* block.

The QSS of the three o-16S rRNA promoter states are: (i) the open free promoter (*g̅*_*r*_), (ii) the promoter when bound by *σ* being actively transcribed (*x̅*_*r*_, calculated using *x̂*_*r*_), or (iii) the promoter bound by the regulator and therefore inhibited (*K̅*_*r*_):

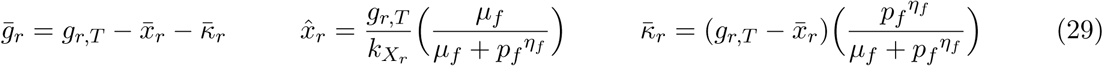

*x̅*_*r*_ determines the rate of host ribosome co-option, via the o-16S rRNA (*r*, see Equation 20) with *g*_*r*, *T*_/*k*_*X*_*r*__ determining the maximal rate (*r*_0_ block, Figure 1b) and *μ*_*f*_/(*μ*_*f*_ + *p*_*f*_^*ηf*^) representing the inhibitory action of the controller *p*_*f*_ (*A* block, Figure 1b).

The rate of change of the orthogonal 16S rRNA is as described in Equation 20 and co-option of the host ribosomes is described in Equation 27.

Finally, the number of free orthogonal ribosomes is given by:

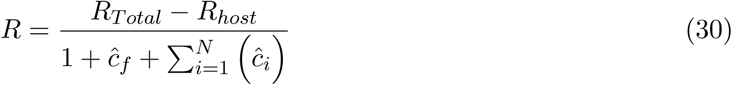

Note that this follows the same form as Equation 27, with the total number of o-ribosomes available to the circuit being the total number of all ribosomes (*R*_*Total*_) minus the number of host ribosomes (*R*_*host*_).

Using the specific binding and unbinding rates of cellular components reported in [17], and calculating their respective dissociation constants as needed, we can compare the behaviour of the full mechanistic model and reduced model. Simulations demonstrate that the reduced model accurately captures the transient and steady-state behaviour of the full model, for both simple circuits based on activation of multiple genes and more complex circuits including oscillatory inputs (Figure 1). Crucially, the model reduction process preserves the rapidly changing closed-loop dynamics produced by the non-linear controller (Figure 1c) The model could potentially be simplified further by making the additional assumption that the equations describing the dynamics of the o-16S rRNA (Equation 20) and host ribosomes (Equation 21) are at steady state, producing a model which tracks only the protein dynamics - whose control is the main subject of this paper. However, this model no longer captures the transient dynamics of the system, although it does still successfully recapitulate the steady state behaviour of the full model to static inputs (Figure S1). In the presence of oscillatory inputs, this additional reduction acts to hide the induction of oscillations in other genes due to the sharing of cellular resources. Analysis of the parameters shows that o-ribosome assembly is slow (*ϱ*_*f*_ = 0.9 (nM·h)^‒1^) violating the assumption that these species are at quasi-steady state.

### 2.2 Model analysis reveals a trade off between gene expression and level of decoupling

Using our model as a design tool, we carried out a multiobjective optimisation of the experimentally tunable controller parameters aiming to produce both high protein levels and low gene coupling (see Methods). We assess the impact on one gene *p*_1_ as a second gene *p*_2_ is induced by a new input *u*_2_ (as described in Figure 3). The difference in expression of *p*_1_ due to the *u*_2_ disturbance is termed ‘coupling’. We assess the impact on protein levels by comparing final protein outputs to those of the same circuit where translation is uncontrolled and mediated by the host ribosome pool. This identified a hard trade-off between these two objectives, with the range of equally optimal solutions (the Pareto-optimal front) showing an inverted concave shape, i.e. decreases in gene coupling are achieved at the expense of decreases in gene expression. Our simulations suggest that coupling can be halved for only a 20% reduction in gene expression, while coupling can be reduced to between 5-10% for a 50% reduction in gene expression (Figure 2).

**Figure 2:**
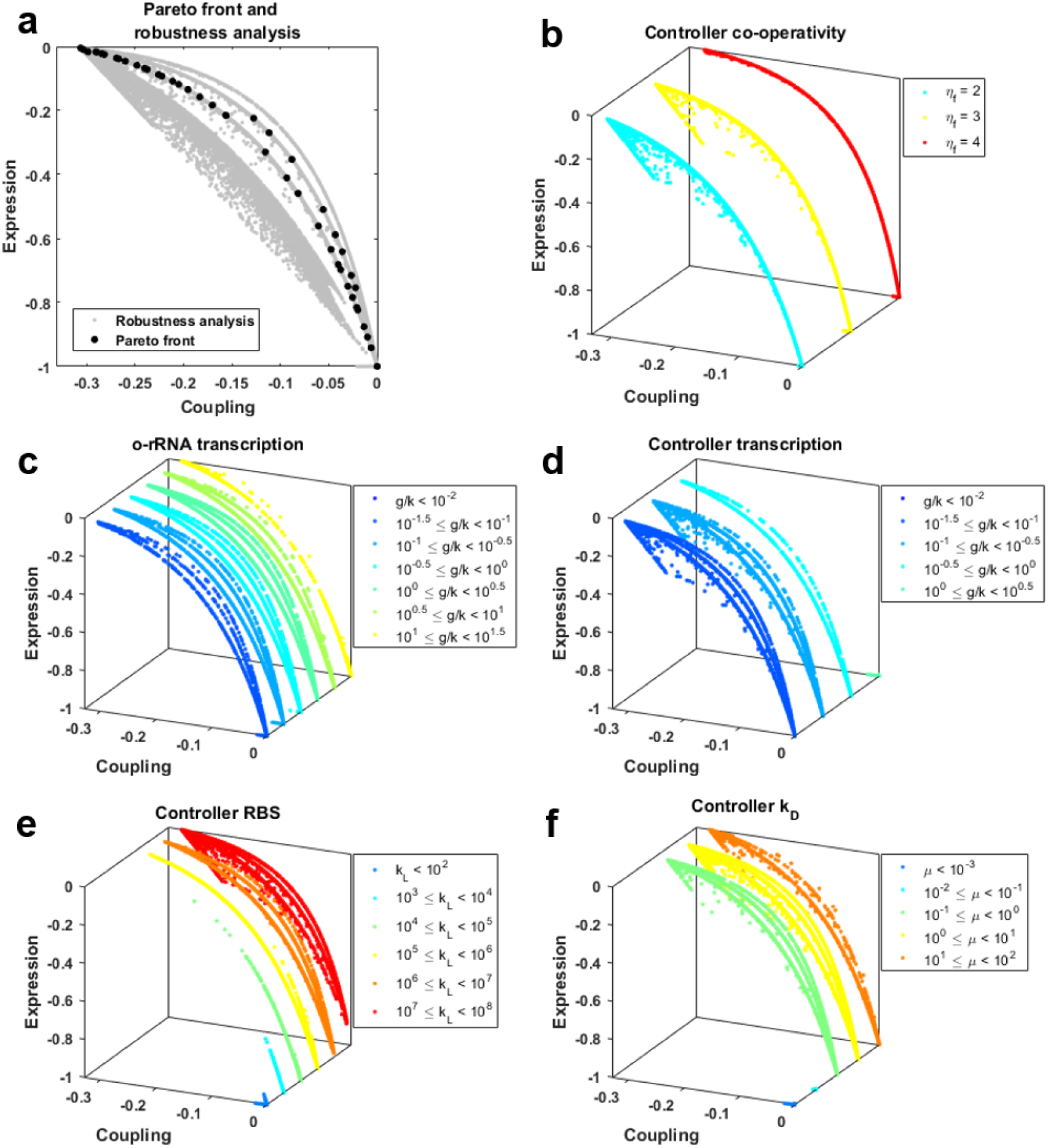
Trade-off between gene expression and decoupling. **(a)** The Pareto front demonstrating the trade off between gene expression and coupling. Expression is measured as change in steady state gene expression in comparison to simulations of the circuit using the saturated host ribosome pool. Coupling is measured as the steady-state change in *p*_1_ in response to *u*_2_. Robustness was determined by allowing the controller parameters to vary within +/-50%. *N* = 89, 890. The values of the respective parameters are shown in the following panels. As described in the main text, controllers where *η*_*f*_ = 1 are removed from the following panels for clarity. Also note that the third axis and subsequent separation serves only to aid visualisation and does not represent parameter value which is indicated by the colour and outlined in the figure legend. **(b)** Controller co-operativity as shown by *η*_*f*_. **(c)** o-rRNA transcription as determined by the *g*_*r*, *T*_/*k*_*X*_*r*__ ratio. **(d)** Transcription of the controller protein as determined by the *g*_*f*, *T*_/*k*_*X*_*f*__ ratio. **(e)** Controller mRNA ribosome binding site strength as measure by mRNA-ribosome dissociation constant *k*_*L*_*f*__. **(f)** Controller protein *g*_*r*_ dissociation constant *μ*_*f*_.

**Figure 3:**
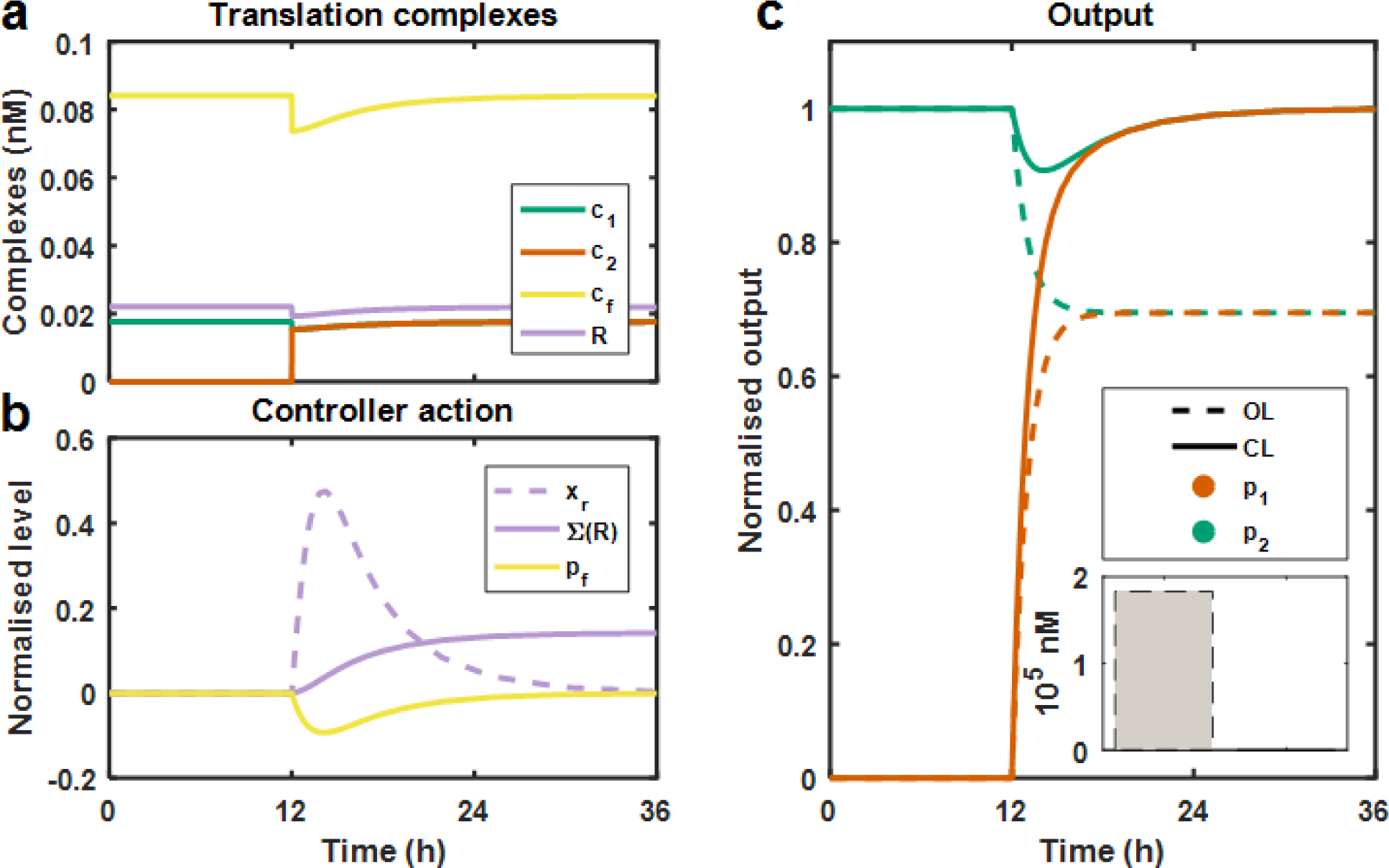
Controller dynamics as it decouples circuit genes. Simulations showing the dynamics of a highly decoupling controller (Point 47, Table S1). The first gene *p*_1_ is constitutively expressed, *u*_1_ = 500 nM throughout. At 12 h, *u*_2_ rises from 0 to 500 nM. **(a)** Translational complexes: Changing the distribution of the orthogonal ribosomes across circuit and contoller mRNAs, *c*_1_, *c*_2_ and *C*_*f*_ represent the translation complexes of the mRNAs of gene 1, 2 and regulator *f* respectively. *R* represents the free orthogonal ribosomes. *C*_*f*_ acts as the sensor for the disturbance at *t* = 12 h. Levels are a normalised by the total number of orthogonal ribosomes at *t* = 12 h. **(b)** Controller action: Changes in controller components over time. Levels are normalised by value at *t* = 12 such that 0 indicates no change. *x*_*r*_, o-16S rRNA gene in the transcribing state; ∑(*R*), number of orthogonal ribosomes; *P*_*f*_, controller protein. **(c)** Normalised protein output. Protein levels are normalised by *p*_1_(*t* = 12) and shown in the absence of the controller (OL, *g*_*f*, *T*_ = 0 nM) and the presence of the controller (CL). Inset, steady state protein levels.

To determine the robustness of the Pareto front to parameter selection, we varied the optimised parameters by up to 50% for each point on the front. None of these designs result in controller failure where expression is lost but coupling is not abolished - i.e. no designs fall into the lower left quadrant. This suggests that potential parameter variations when selecting biological components to implement the controller can act to move the controller along the front, but should *not* result in failure. We did find that a small number of these perturbed designs show slower responses and we discount these from further analysis. We carried out an additional robustness analysis allowing *all* parameters governing the controller behaviour to vary. This includes parameters which are either difficult to design (e.g. controller translation rate *γ*_*f*_) or intrinsic properties which cannot be designed (e.g. o-rRNA association rate, *μ*_*r*_). We find that all of these controllers also fall upon the same front demonstrating that uncertainty in these values does not preclude controller design (Figure S2).

To determine how each parameter contributes to the gene expression and coupling trade off, we analysed how each changes across the front. This highlights the need for high *η*_*f*_ values. This parameter represents the level of co-operativity in the system brought about for example by transcription factor dimerisation or the presents of multiple operator sites. The true Pareto front coincides with a value of *η*_*f*_ = 4 (Figure 2b). For this reason we discount controllers where *η*_*f*_ = 1 from further analysis in this section as these controllers perform most poorly. We also find that small *μ*_*f*_ values are most often associated with controllers which act to nearly completely decouple genes but at a significant cost to gene expression (Figure 2f). Similarly, small *k*_*L*_*f*__ values, corresponding to strong ribosomes binding sites (low ribosome-mRNA dissociation constant), are associated with large levels of decoupling at a high cost to gene expression (Figure 2e). Simulations suggest *k*_*L*_*f*__ > 10^5^ nM and *μ*_*f*_ > 0.1nM in all cases, for the simple two gene circuit example used here. (Note that for many natural transcription factors co-opted into synthetic gene networks *μ*_*f*_ < 0.1 nM and *η*_*f*_ may be limited. We demonstrate how this can be compensated for in controller design in Section 2.5). A high *g*_*f*, *T*_/*k*_*x*_*f*__ ratio (*g*_*f*, *T*_/*k*_*x*_*f*__ > 1, produced by expressing the regulator from a strong promoter carried on a high copy number plasmid) results in complete decoupling and abolition of gene expression (Figure 2d). We therefore suggest keeping *g*_*f*, *T*_/*k*_*X*_*f*__ < 1 in all instances. We find that the *g*_*r*, *T*_/*k*_*X*_*r*__ ratio governing maximal o-rRNA transcription rate varies significantly across all behaviours making general guidelines difficult to establish (Figure 2c).

To provide specific quantitative design rules, we select five points across the front and assess the parameter space around these points which give rise to these controller behaviours (Table 1). Assessing how parameters correlate in these local clusters shows that *k*_*L*_*f*__ is a key regulator of behaviour. *k*_*L*_*f*__ is inversely correlated with *g*_*r*, *T*_/*k*_*X*_*r*__, indicating that as the o-ribosome production rate increases, a stronger RBS is needed for controller function (i.e. a smaller value of *k*_*L*_*f*__). We also identify an inverse correlation between *k*_*L*_*f*__ and *μ*_*f*_ in the cluster around (−0.01, −0.9), i.e. the most decoupled parameter set, such that decreases in repression by the transcription factor (*μ*_*f*_) can be compensated for by increasing the RBS strength (decreasing *k*_*L*_*f*__). We also find that changes in *k*_*L*_*f*__ and *g*_*f*, *T*_/*k*_*X*_*f*__ have some compensatory effects such that increases in *k*_*L*_*F*__ (i.e. weakening the RBS) can be mitigated by increasing *g*_*f*, *T*_/*k*_*X*_*f*__ (e.g. by increasing copy number).

**Table 1:**
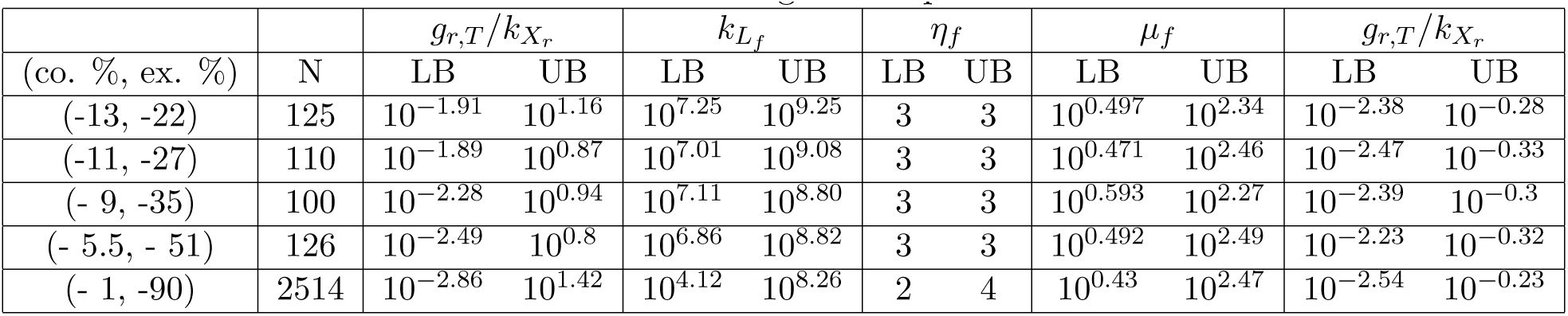
Controller designs to manage the coupling expression trade-off. Regions of parameter space were identified as described in the Methods using a distance score of 0.25. The coupling (co. %) and expression cost (ex. %) are reported for each controller from the Pareto front chosen. The number of controllers in the local region is reported as *N*.

| | | $g_{r,T}/k_{X_r}$ | | $k_{L_f}$ | | $\eta_f$ | | $\mu_f$ | | $g_{r,T}/k_{X_r}$ | |
| --- | --- | --- | --- | --- | --- | --- | --- | --- | --- | --- | --- |
| (co. %, ex. %) | N | LB | UB | LB | UB | LB | UB | LB | UB | LB | UB |
| (-13, -22) | 125 | $10^{-1.91}$ | $10^{1.16}$ | $10^{7.25}$ | $10^{9.25}$ | 3 | 3 | $10^{0.497}$ | $10^{2.34}$ | $10^{-2.38}$ | $10^{-0.28}$ |
| (-11, -27) | 110 | $10^{-1.89}$ | $10^{0.87}$ | $10^{7.01}$ | $10^{9.08}$ | 3 | 3 | $10^{0.471}$ | $10^{2.46}$ | $10^{-2.47}$ | $10^{-0.33}$ |
| (- 9, -35) | 100 | $10^{-2.28}$ | $10^{0.94}$ | $10^{7.11}$ | $10^{8.80}$ | 3 | 3 | $10^{0.593}$ | $10^{2.27}$ | $10^{-2.39}$ | $10^{-0.3}$ |
| (- 5.5, - 51) | 126 | $10^{-2.49}$ | $10^{0.8}$ | $10^{6.86}$ | $10^{8.82}$ | 3 | 3 | $10^{0.492}$ | $10^{2.49}$ | $10^{-2.23}$ | $10^{-0.32}$ |
| (- 1, -90) | 2514 | $10^{-2.86}$ | $10^{1.42}$ | $10^{4.12}$ | $10^{8.26}$ | 2 | 4 | $10^{0.43}$ | $10^{2.47}$ | $10^{-2.54}$ | $10^{-0.23}$ |

We demonstrate the functioning of the controller using the design with the lowest level of gene coupling, point 47 on the Pareto-optimal front (Table S1). The equivalent analysis for the intermediate point 9 is shown in Figure S3). The controller successfully insulates one gene from the induction of another (Figure 3), bar a short transient disturbance (<12 h). Tracking the concentrations of the intermediate species reveals the operation of the controller; with translation of *p*_*f*_ falling (Figure 3b) and the number of o-rRNA genes being transcribed increasing as the second gene is induced (Figure 3b). This results in a net increase in the number of orthogonal ribosomes (Figure 3b) which means that in the long term the translation complexes producing each protein do not change (Figure 3a).

### 2.3 Designing system response times by tuning controller parameters

Analysis of the controllers tested so far has focused on how they are able to correct steady state errors brought about by gene coupling and so we have largely ignored the system dynamics, bar excluding excessively slow controllers (e.g. penalising simulations which only reach steady state after > 24 h). However, a controller which decouples genes well but has a slow response time will not be suitable for many applications in synthetic biology. Therefore we took the previous candidate controllers and conducted a local sensitivity analysis around each design point to assess the impact of each parameter on the system’s speed of response. In addition to the controller parameters varied so far we also varied *δ*_*ρ*_, *δ*_*m*_*f*__ and *δ*_*P*_*f*__, which represent the decay of the o-rRNA, controller mRNA and controller protein respectively. These parameters were kept constant in the previous design evaluations to minimise the number of parameters in the optimisation, but since decay parameters often have significant affects on speed of response we explicitly assess their impact here.

Beginning with the four controllers which show intermediate behaviours (i.e. do not show complete decoupling), changing the parameters has significant impact on coupling and expression levels, as discussed above. For controllers which have strong decoupling ability we find that there are individual parameters which when varied do not significantly affect the decoupling ability (e.g. *g*_*r*, *T*_/*k*_*X*_*r*__, *μ*_*f*_) although these do still affect the expression levels (i.e. the controller gain).

In all five cases, the o-rRNA decay constant *δ*_*r*_) and protein controller decay constant (*δ*_*p*_*f*__) are key to determining the speed of the system response. Increasing both parameters acts to increase the speed of response, at an expense of decoupling ability (Figure 4). In the regions tested, varying *δ*_*r*_. is less likely to introduce significant overshoots into the system (as seen at low *δ*_*P*_*f*__ values). However, a greater range of speed-up is achievable by varying the protein decay constant. The latter is also a more experimentally tractable parameter. Increasing both parameters acts antagonistically, with increases in *δ*_*r*_. decreasing gene coupling and increases in *δ*_*p*_*f*__ increasing it, meaning tuning both parameters may be advantageous. We see very little impact from varying the mRNA decay rate *δ*_*m*_*f*__).

**Figure 4:**
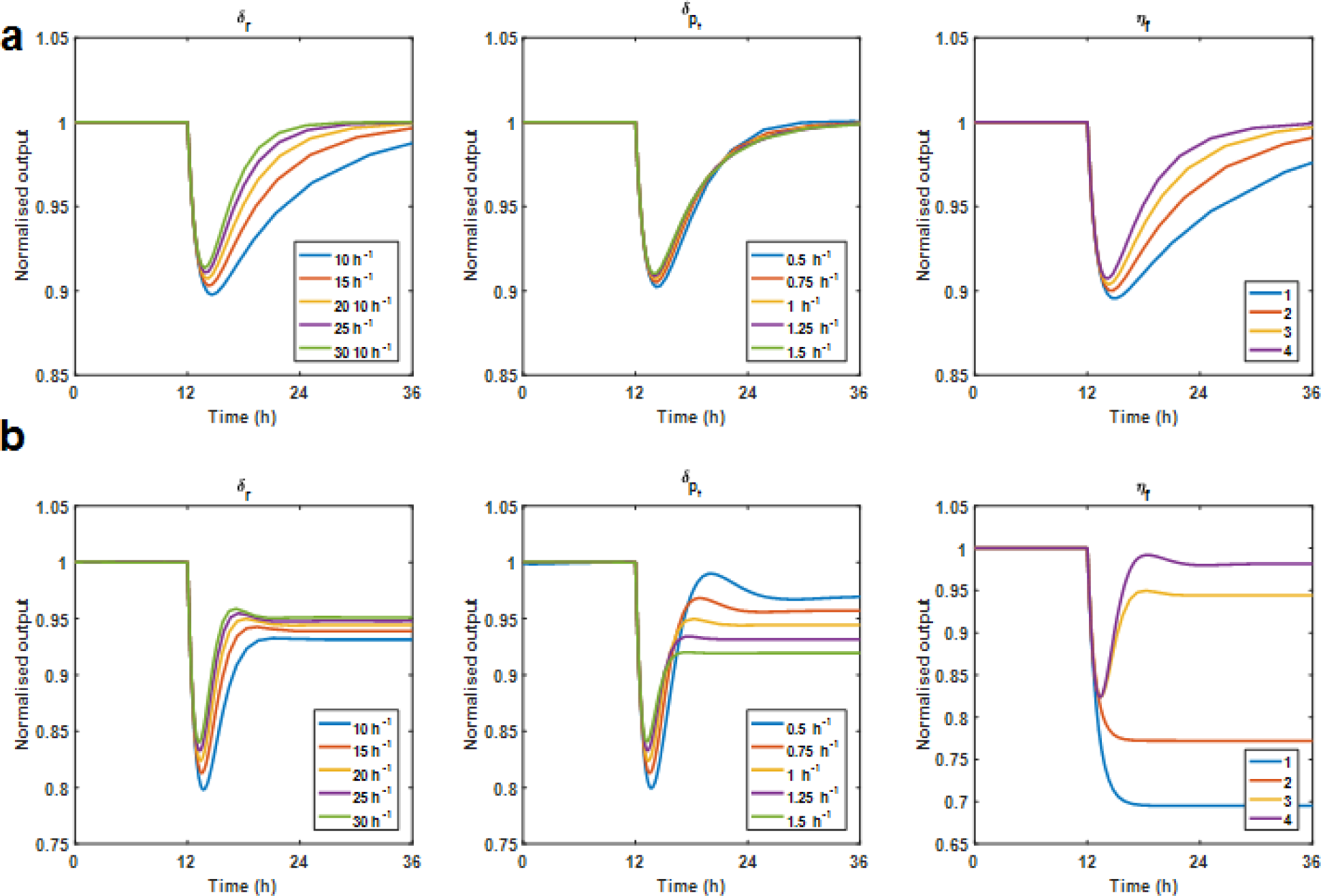
Tuning decay parameters allows design of system dynamics. The effect of varying the decay prameters *δ*_*r*_ and *δ*_*P*_*f*__ on the response of *p*_1_ to the additional input *u*_2_ (as described in Figure 3. The optimal parameter as determined by the optimisation routine was varied by +/ – 25% and +/ – 50% named *δ* in the figure legend. **(a)** Sensitivity analysis around the parameter set from the high decoupling regime (Point 47). **(b)** Sensitivity analysis around a parameter set from the intermediate decoupling regime (Point 9).

As previously discussed the value of the controller co-operativity *ηf*) is a key determinant of controller decoupling ability (Figure 4). This analysis replicates this result and also highlights that, at least in this parameter regime, increasing co-operativity also acts to significantly increase the speed of response.

### 2.4 Potential biological implementations of the controller designs

We carried out a detailed literature review to identify potentially suitable repressors with which to implement our system, focusing our analysis on (i) the ability of the repressor to be expressed in bacterial hosts (i.e. repressors from bacteria or bacteriophage), (ii) orthogonality (i.e. repressors which are not used in fundamental host processes), (iii) the presence of a known promoter architecture (which could be used to infer the dissociation constant of the RNA polymerase, see Section S2) and (iv) detailed characterisation of binding kinetics (ideally dissociation constants measured in a biochemical assay, rather than a constant inferred from device function such as by fitting a Hill function to induction-fluorescence curves, as is often the case). We identified six repressors from this literature search, including the commonly used LacI [18], TetR [19] and cI [20] repressors. We also identified putative controller candidates Cro and RstR from bacteriophages PY54 [21], CTX*φ* [22] and LmrR, a global regulator of antibiotic resistance from Gram positive *Lactococcus lactis* [23].

Using the results of our sensitivity analysis and additional biological constraints we identified a number of feasible biological implementations; (i) the o-rRNA and regulator having the same medium copy number (mimicking placement in the same plasmid, such as ColE1), (ii) a high copy number regulator, carried on for example a pUC vector, and a chromosomally integrated regulator and (iii) the effect of a protein degradation motif on designs of type (ii). Note that we did not assess the potential designs requiring the o-rRNA and regulator to be carried on different copy number plasmids, as these would result in high burden on the cells and significantly decreased growth rate as these cells would need to carry at least three plasmids, one containing circuit genes and one each for the o-rRNA gene and regulator. All of these prototype controllers fall along the Pareto front (Figure 5a).

**Figure 5:**
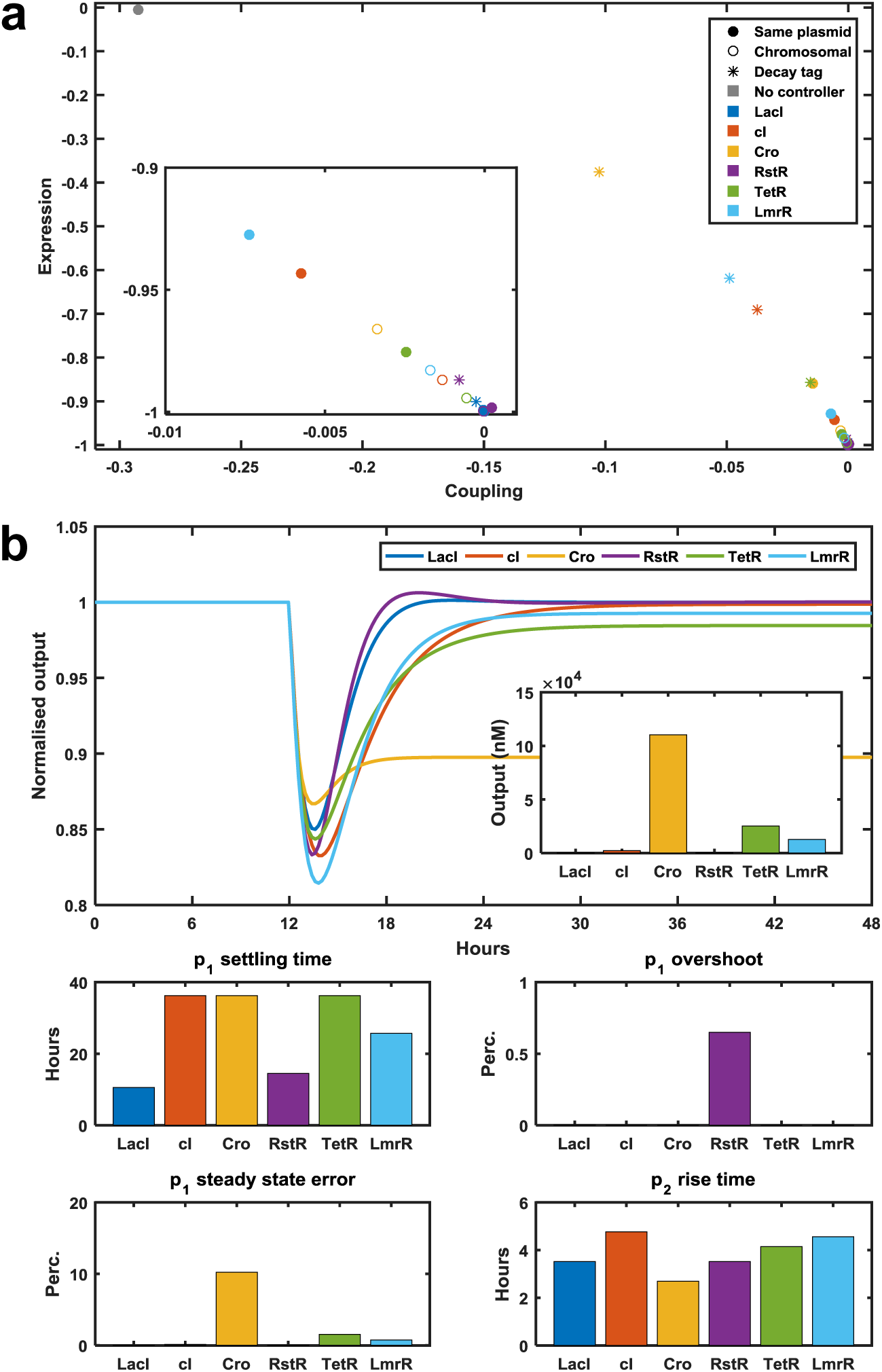
Comparison of biological implementations based on orthogonal repressors. Simulations of implementations based on the repressors in different plasmid confirmations and with degradation motifs. **(a)** The positions of the prototype controllers in the coupling-expression space. Inset, expansion of the main figure around point (0,-1). Point colours represent the regulator protein and point style denotes copy number as follows: Same plasmid, *g*_*r*, *T*_ = *g*_*f*, *T*_ = 100 nM; Chromosomal, *g*_*r*, *T*_ = 500 nM and *g*_*f*, *T*_ = 10 nM. Decay tag, *g*_*r*, *T*_ = 500 nM, *g*_*f*, *T*_ = 10 nM, *δ*_*p*_*f*__ = 15 h^‒1^ ≈ *t*_1_/_2_ = 3 – 5 minutes, equivalent to LVA tag []. **(b)** *Upper* Circuit dynamics showing the normalised levels of protein 1. Inset, steady state output at *t* = 48 h. *Lower* Characterisation of the response of *p*_1_ to the disturbance caused by *u*_2_. Settling time, number of hours from the induction until *p*_1_ returns to steady state; Overshoot, transient increase in *p*_1_. Steady state error, difference between *p*_1_(*t* = 48) and *p*_1_(*t* = 12). Rise time, time it takes for *p*_2_ to increase from 10% of its steady state to 90% of its steady state. Designs are available in Table S4.

We carried forward designs representing a range of trade-offs between gene expression and decoupling for further analysis (Table S4). The putative controllers based on tetramers (LacI and RstR) show the fastest dynamics and best decoupling with minimal *p*_1_ settling times and *p*_2_ rise time upon induction of the second gene (Figure 5b). Designs based on the phage repressor Cro with degradation motifs show the highest gene expression with an acceptable *p*_2_ rise time of < 3 hours but coupling is far from fully abolished (Figure 5b).

### 2.5 A dynamic resource allocation controller restores modularity in a range of more complex gene circuits

Having successfully demonstrated the ability of the proposed approach to decouple two independent modules, we analyse the ability of the controller to remove resource dependent failure in a variety of more complex gene circuits (Figure 6).

**Figure 6:**
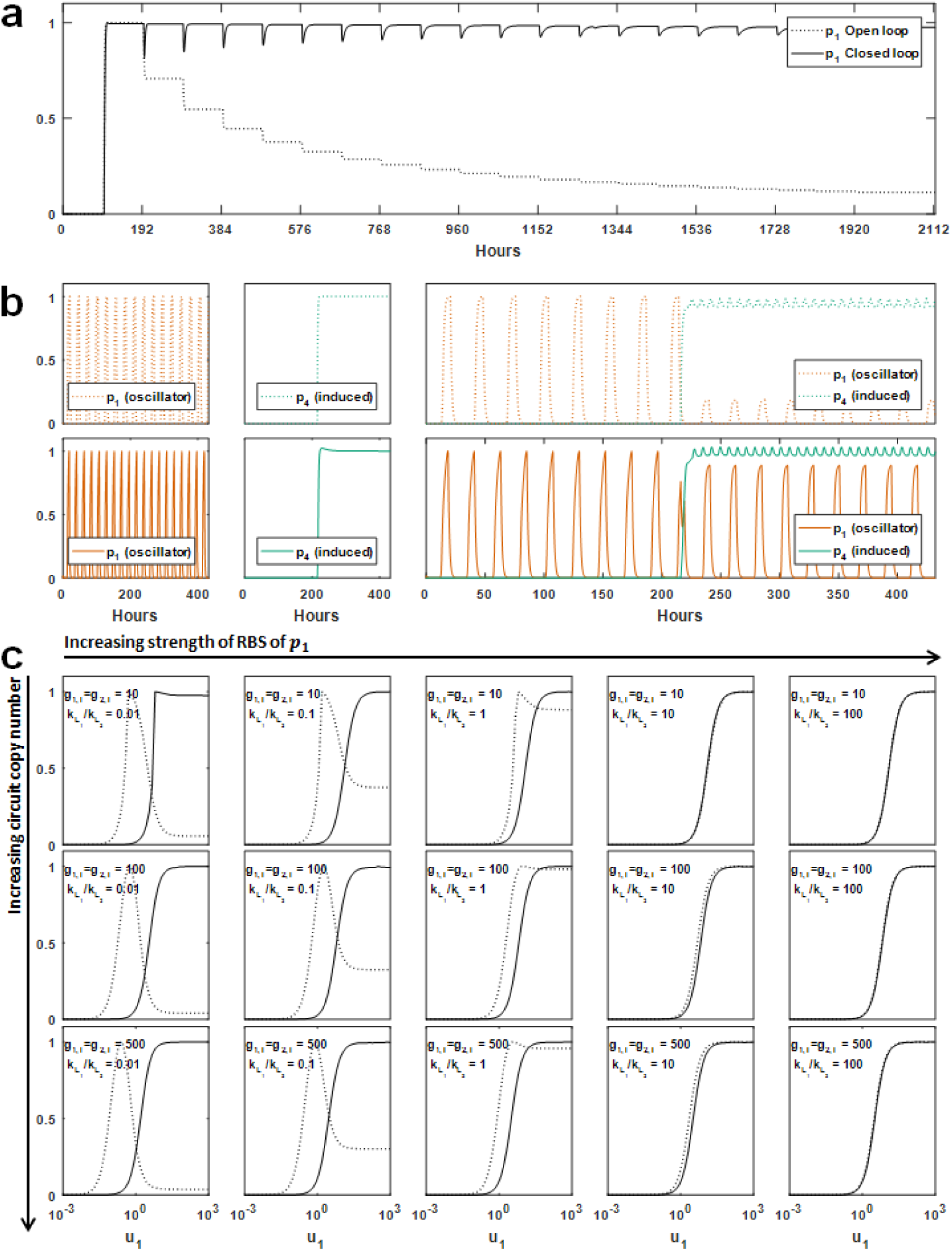
The controller rescues modularity in a variety of circuit contexts. A range of common circuits were simulated in both the open and closed loop confirmations. All y-axes are normalised output. **(a)** The controller successfully renders a gene invulnerable to the induction of many additional genes at 100 h intervals. Other genes not shown.**(b)** Maintaining repressilator behaviour in the presents of an induced gene. The repressilator (protein *p*_1_ to *p*_3_, only *p*_1_ is shown) is simulated before an additional gene *p*_4_ with a stronger RBS is induced. *Upper panels* Open loop (no controller). *Lower panels* Closed loop (with controller). *Left and centre panels* Function of the individual modules alone. *Right* Function of the two modules in one circuit. *p*_4_ is induced at 24 h. **(c)** The controller removes resource limitation-induced failure in the design of an activation cascade (*u*_1_ → *p*_1_ → *p*_2_). In the absence of the controller (dotted line) some prototype designs do not show the monotonically increasing output of *p*_2_ to *u*_1_ as desired in an activation cascade. The controller removes these resource limitations allowing the circuit to function as expected across all prototype designs.

We initially simulate multiple SISO modules with new modules being activated at different intervals. In the absence of the controller, activation of each additional module has an impact on the previously activated modules. For example, the expression of the first module *p*_1_ falls by over 50% as three additional genes are induced. As shown in Figure 6a, the controller successfully eliminates this coupling, making *p*_1_ relatively insensitive to the induction of over 10 additional genes (although note that the rise time and settling time increase slightly with the induction of each additional gene).

A key aim of synthetic biology is that previously characterised components or devices can be introduced into the same cell to form a complex circuit. Here we assess the effect of introducing two separately characterised devices into one complex circuit, i.e. we want to investigate what is the effect of introducing an additional resource consumer on a previously characterised device. As the production of robust genetic oscillators to create clocks for temporal processes functions is of fundamental importance in synthetic circuit design, we consider designs for the repressilator clock and an additional SISO module. These modules are first simulated separately, as shown in (Figure 6b, *upper panel*). Upon linking these separate devices through a common pool of resources, i.e. coupled through their competition for ribosomes, we see that *p*_4_ induction destroys the oscillations of the repressilator (Figure 6b, upper right panel). If, however, we consider the design of these two devices in the presence of the controller and then introduce them into the same resource pool as before, we see that circuit function is now maintained (Figure 6b, *lower panels*). Note that while there is still a small loss in repressilator amplitude upon induction of *p*_4_ this is significantly reduced, thus staying closer to the original device behaviour.

It has previously been shown that resource limitations can change the input-output response of a simple genetic activation cascade [4]. The authors show that if the upstream module has a stronger ability to sequester ribosomes than the downstream module (a small *k*_*L*_1__-to-*k*_*L*_2__ ratio) then the expected response determined from simple Hill-function type modelling (i.e. an increasing output to increasing input in a step-like fashion) can become biphasic or even invert (Figure 6c, dotted open loop lines). We simulate a range of prototype activation cascades in the absence and presence of our controller. In the absence of the controller, no additional resources are available as demand increases and so we see the activation cascade failing in the same manner as found in Qian *et al.*, In the presence of the controller, the desired behaviour of the activation cascade is restored, as translational capacity is directed to the circuit as demand increases. The controller acts to remove the resource limitation, thus allowing simpler models, which often do not account for limited cellular resources, to be used to produce circuit designs which then function as expected *in vivo.*

### 2.6 Conclusions

Numerous genetic components and devices have been developed to ensure predictable gene expression or dampen the effect of loading in genetic circuits. However, to date, little attention has been paid to developing genetic devices that are capable of relieving cellular resource limitations. Controllers for orthogonal transcriptional activity based on phage RNA polymerases have been developed [24, 25] and we have previously implemented a prototype translational controller [14]. Here, we develop a detailed mechanistic model of gene expression and resource allocation, which when simplified to a tractable level of complexity, allows the rational design of optimal translational controllers. We demonstrated that this new model allows the design of controllers which can dynamically allocate orthogonal ribosomes to synthetic circuits within reasonable timeframes (< 12 hours). Using our model, we identify a fundamental trade-off in controller design; that reducing coupling act to decrease gene expression. We determined how each controller design parameter affects the overall closed-loop behaviour of the system, leading to a detailed set of design guidelines for optimally managing this trade-off. We find that both controller co-operativity and RBS strength are key parameters in determining the level of decoupling that can be achieved. Based on our designs, we identified and evaluated a number of alternative potential experimental implementations of the proposed system using commonly available biological components. Finally, we showed that our controller is capable of dynamically allocating ribosomes as needed to restore modularity in a number of more complex synthetic circuits, such as the repressilator, and activation cascades composed of multiple interacting modules.

## 3 Methods

### 3.1 Numerical simulations

All models were implemented in either MATLAB 2016b and 2017a (The MathWorks Inc, MA, USA) and simulated using the in-built stiff solvers *ode15s* and *ode23s* using increased tolerances (Relative 10^‒6^ and Absolute 10^‒6^). Simulations were deemed to have reached steady state when the maximum of the calculated derivative was less than 10^‒2^ (relaxed) or 10^‒3^ (strict). Additional specialist functions as needed were utilised from the Optimization Toolbox (Version 7.4) and Parallel Computing Toolbox (Version 6.8 or 6.10).

### 3.2 Assessment of controller function

The behaviour of controllers was characterised by simulating the action of a simple two gene circuit. Initially, the behaviour of one gene *p*_1_ is simulated before its response is assessed to the induction of a second gene *p*_2_ at time *t* = *θ*_*ind*_. Coupling and expression are normalised:

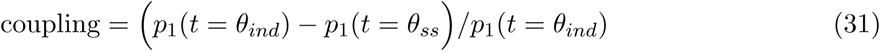

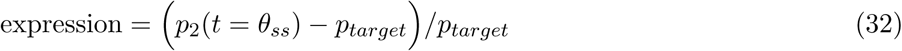

### 3.3 Optimisation

The mutliobjective optimisation was carried out using the inbuilt function *gamuliobj* with a population size of 200 individuals and with a Pareto fraction of 0.25 from the Optimization Toolbox. *k*_*X*_ values were set to 1 allowing the *x̂* ratios to be investigated by varying *g*_*r*_*T*__ and *g*_*f*_,_*T*_ only. See Section S2 for a discussion of permissible parameter bounds. The parameters varied (and their scale and bounds) were *g*_*r*, *T*_/*k*_*X*_*r*__ ratio (log10 scale, 10^‒2^ – 10^2^), *k*_*L*_*f*__ (log10 scale, 10^3^ – 10^8^), *η*_*f*_ (linear scale, 1 – 4), *μ*_*f*_ (log10 scale, 10^‒2^ – 10^3^) and *g*_*r*, *T*_/*k*_*X*_*r*__ ratio (log10 scale, 10^‒2^ – 10^2^). The optimisation routine aims to minimise:

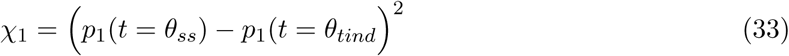

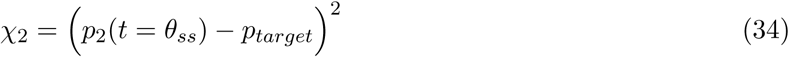

where *θ*_*ind*_ is the time of the induction of *p*_2_ and *θ*_*ss*_ is the last time point, *t*_*max*_. If the simulation is not at steady state at *t*_*max*_ then the result is given the poorest fitness. *p*_*target*_ is calculated by simulating the action of the circuit in a model utilising the host ribosome pool for gene expression.

### 3.4 Selection of controller parameters for design guidelines

Coupling and expression scores where calculated for each controller as outlined above for all the results of robustness analysis. These results were then scaled by their maximum absolute values to ensure both axes are between 0 and 1 (note that for calculating the distance metric we can ignore signs). We calculate the Euclidean distance between each point (*x*_*s*_*caled*, *y*_*s*_*caled*) and the point of interest from our numerical optimisation (*x*_0_,*y*_0_):

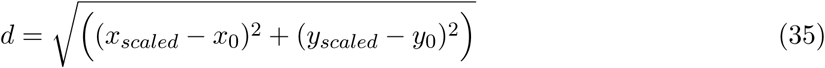

We then define any points within a circuit of radius 0.025 centred on (*x*_0_, *y*_0_) as within the local region. Qualitatively these points have the same behaviour and so we group them for further analysis as outlined in the main text.

## References

[1] J. A. Brophy and C. A. Voigt, “Principles of genetic circuit design.,” Nature Methods, vol. 11, no. 5, pp. 508–20, 2014.

[2] S. Cardinale and A. P. Arkin, “Contextualizing context for synthetic biology - identifying causes of failure of synthetic biological systems,” Biotechnology Journal, vol. 7, no. 7, pp. 856–866, 2012.

[3] D. Del Vecchio, A. J. Ninfa, and E. D. Sontag, “Modular cell biology: retroactivity and insu-lation.,” Molecular Systems Biology, vol. 4, no. 161, p. 161, 2008.

[4] Y. Qian, H.-H. Huang, J. Jimenez, and D. Del Vecchio, “Resource competition shapes the response of genetic circuits,” ACS Synthetic Biology, vol. 6, no. 7, pp. 1263–1272, 2017.

[5] C. Lou, B. Stanton, Y.-J. Chen, B. Munsky, and C. A. Voigt, “Ribozyme-based insulator parts buffer synthetic circuits from genetic context.,” Nature biotechnology, vol. 30, no. 11, pp. 1137–42, 2012.

[6] C. J. Bashor and J. J. Collins, “Insulating gene circuits from context by RNA processing,” Nature Publishing Group, vol. 30, no. 11, pp. 1061–1062, 2012.

[7] K. S. Nilgiriwala, J. Jimenez, P. M. Rivera, D. D. Vecchio, J. Jimenez, P. M. Rivera, and D. Del Vecchio, “Synthetic tunable amplifying buffer circuit in E. coli.,” ACS synthetic biology, vol. 4, pp. 577–84, 2015.

[8] M. Scott, C. W. Gunderson, E. M. Mateescu, Z. Zhang, and T. Hwa, “Interdependence of cell growth and gene expression: origins and consequences.,” Science, vol. 330, no. 6007, pp. 1099–1102, 2010.

[9] F. Ceroni, R. Algar, G.-B. Stan, and T. Ellis, “Quantifying cellular capacity identifies gene expression designs with reduced burden,” Nature Methods, vol. 12, no. 5, pp. 415–423, 2015.

[10] A. Gyorgy, J. I. Jimenez, J. Yazbek, H.-H. Huang, H. Chung, R. Weiss, and D. Del Vecchio, “Isocost Lines Describe the Cellular Economy of Genetic Circuits,” Biophysical Journal, vol. 109, no. 3, pp. 639–646, 2015.

[11] M. Carbonell-Ballestero, E. Garcia-Ramallo, R. Montanez, C. Rodriguez-Caso, and J. Marcia, “Dealing with the genetic load in bacterial synthetic biology circuits : convergences with the Ohm’s law,” Nucleic Acids Research, vol. 44, no. 1, pp. 496–507, 2016.

[12] T. E. Gorochowski, I. Avcilar-Kucukgoze, R. A. Bovenberg, J. A. Roubos, and Z. Ignatova, “A minimal model of ribosome allocation dynamics captures trade-offs in expression between endogenous and synthetic genes,” ACS Synthetic Biology, vol. 5, pp. 710–720, 2016.

[13] T. Shopera, L. He, T. Oyetunde, Y. J. Tang, and T. S. Moon, “Decoupling resource-coupled gene expression in living cells,” ACS Synthetic Biology, vol. 6, no. 8, pp. 1596–1604, 2017.

[14] A. P. S. Darlington, J. Kim, J. I. Jimenez, and D. G. Bates, “Dynamic Allocation Of Orthogonal Ribosomes Facilitates Uncoupling Of Co-Expressed Genes,” Accepted by Nature Communications. Pre-print available at https://doi.org/10.1101/138362, 2017.

[15] R. Milo and R. Phillips, Cell Biology by the numbers. New York, NY: Garland Science, 2016.

[16] H. Bremer, P. Dennis, and M. Ehrenberg, “Free RNA polymerase and modeling global transcription in Escherichia coli,” Biochimie, vol. 85, no. 6, pp. 597–609, 2003.

[17] A. Hamadeh and D. del Vecchio, “Mitigation of resource competition in synthetic genetic circuits through feedback regulation,” Proc. of 53rd IEEE Conference on Decision and Control, pp. 3829–3834, 2015.

[18] C. M. Falcon and K. S. Matthews, “Operator DNA sequence variation enhances high affinity binding by hinge helix mutants of lactose repressor protein,” Biochemistry, vol. 39, no. 36, pp. 11074–11083, 2000.

[19] A. Kamionka, J. Bogdanska-Urbaniak, O. Scholz, and W. Hillen, “Two mutations in the tetracycline repressor change the inducer anhydrotetracycline to a corepressor,” Nucleic Acids Research, vol. 32, no. 2, pp. 842–847, 2004.

[20] Y. Wang, L. Guo, I. Golding, E. C. Cox, and N. P. Ong, “Quantitative transcription factor binding kinetics at the single-molecule level,” Biophysical Journal, vol. 96, no. 2, pp. 609–620, 2009.

[21] J. A. Hammerl, N. Roschanski, R. Lurz, R. Johne, E. Lanka, and S. Hertwig, “The molecular switch of telomere phages: High binding specificity of the PY54 Cro lytic repressor to a single operator site,” Viruses, vol. 7, no. 6, pp. 2771–2793, 2015.

[22] H. H. Kimsey and M. K. Waldor, “The CTXϕ Repressor RstR Binds DNA Cooperatively to Form Tetrameric Repressor-Operator Complexes,” Journal of Biological Chemistry, vol. 279, no. 4, pp. 2640–2647, 2004.

[23] H. Agustiandari, E. Peeters, J. G. de Wit, D. Charlier, and A. J. M. Driessen, “LmrR-mediated gene regulation of multidrug resistance in Lactococcus lactis,” Microbiology, vol. 157, no. 5, pp. 1519–1530, 2011.

[24] T. H. Segall-Shapiro, A. J. Meyer, A. D. Ellington, E. D. Sontag, and C. A. Voigt, “A ‘resource allocator’ for transcription based on a highly fragmented T7 RNA polymerase.,” Molecular Systems Biology, vol. 10, no. 7, p. 742, 2014.

[25] M. Kushwaha and H. M. Salis, “A portable expression resource for engineering cross-species genetic circuits and pathways,” Nature Communications, vol. 6, p. 7832, 2015.

